# Induced Polarization in MD Simulations of the 5HT_3_ Receptor Channel

**DOI:** 10.1101/2020.03.01.971853

**Authors:** Gianni Klesse, Shanlin Rao, Stephen J. Tucker, Mark S.P. Sansom

## Abstract

Ion channel proteins form water-filled nanoscale pores within lipid bilayers and their properties are dependent on the complex behavior of water in a nano-confined environment. Using the pore of the 5HT3 receptor (5HT3R) we compare additive with polarizable models in describing the behavior of water in nanopores. Molecular Dynamics simulations were performed with four conformations of the channel: two closed state structures, an intermediate state, and an open state, each embedded in a phosphatidylcholine bilayer. Water density profiles revealed that for all water models, the closed and intermediate states exhibited strong dewetting within the central hydrophobic gate region of the pore. However, the open state conformation exhibited varying degrees of hydration, ranging from partial wetting for the TIP4P/2005 water model, to complete wetting for the polarizable AMOEBA14 model. Water dipole moments calculated using polarizable force fields also revealed that water molecules remaining within dewetted sections of the pore resemble gas phase water. Free energy profiles for Na+ and for Cl− ions within the open state pore revealed more rugged energy landscapes using polarizable force fields, and the hydration number profiles of these ions were also sensitive to induced polarization resulting in a substantive reduction of the number of waters within the first hydration shell of Cl− whilst it permeates the pore. These results demonstrate that induced polarization can influence the complex behavior of water and ions within nanoscale pores and provides important new insights into their chemical properties.

**ToC Graphic:** 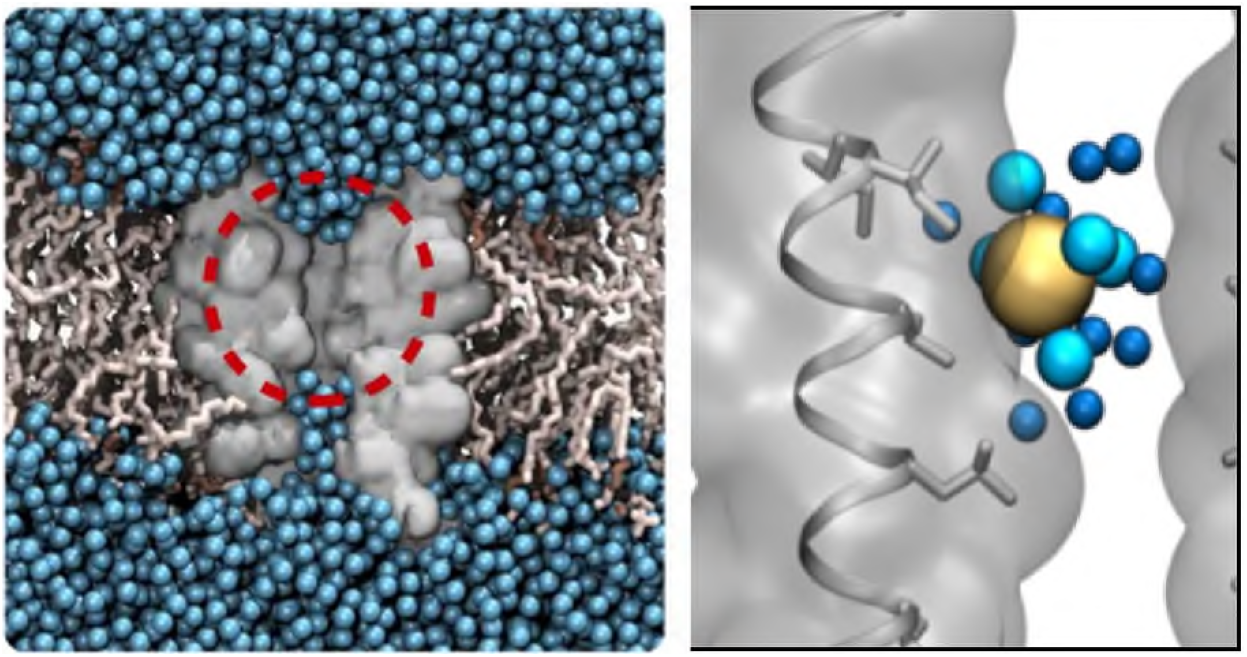

## Introduction

Ion channel proteins form structurally dynamic nanoscale pores within cell membranes ^1–2^ where they are responsible for regulating the movement of ions across the lipid bilayer. Their activity underlies nearly all forms of cellular electrical activity and signaling, and they are an important class of therapeutic targets for the treatment of disease.

When open and conductive, the typical ion channel transmembrane pore has an internal radius of ~0.5 nm and length ~5 nm. They are typically filled with water, thus providing low-energy ion permeation pathways across a membrane. Their functional properties are therefore highly dependent on the complex behavior of water and ions in such nano-confined environments. Molecular simulations play a key role in understanding these anomalous properties ^3–5^. However, the rapidly increasing number of high-resolution structures now available for ion channel pores demand faster and more accurate computational approaches for predicting their functional properties.

The permeation of ions through a sub-nm diameter pore is influenced by both the pore radius and the local hydrophobicity of the pore lining ^6^. Thus, although permeation can occur through polar regions only just larger than the radius of the permeating ion, a hydrophobic pore region of comparable dimensions may undergo spontaneous de-wetting ^7–9^ to present an energetic barrier to permeation without complete steric occlusion. This has been referred to as a hydrophobic gate ^7, 10–12^. The concept of hydrophobic gating has been explored in a number of different channels, in particular several members of the pentameric ligand-gated ion channels (pLGICs), including the nicotinic acetylcholine receptor ^13^, GLIC ^11^, ^14^, and 5-HT_3_ (serotonin) receptor ^15–17^, as well as several different classes of tetrameric cation channels. Building on these studies, we have developed a computational tool (CHAP) capable of predicting the conductive state of a pore ^18^ as well as a heuristic approach based upon CHAP analysis of simulations of water for the current ion channel proteome ^6^.

However, the majority of molecular dynamics (MD) simulations of ion channels, including those on which CHAP is based, employ pairwise additive force fields where electrostatic interactions are modelled as Coulombic forces between fixed point charges on atomic centers. Unfortunately, such force fields do not capture the effect of induced polarization where the electronic structure of an atom is altered by the varying distribution of charges in its immediate environment ^19^. Although additive force fields can include induced polarization in a mean-field fashion ^20^, they cannot accurately describe a molecule that exists in both the gas and condensed phases. This potential problem is particularly relevant to studies using water molecules, where the liquid-state dipole moment (2.95 D) obtained from *ab initio* calculations ^21^ is 60% greater than the gas-phase value (1.85 D) ^22^. Likewise, the molecular dipole moment is also expected to vary depending on whether nearby surfaces within the pore are hydrophobic or hydrophilic.

Such limitations are therefore relevant in studies of hydrophobic gating ^12^ where the equilibrium between the “wet” and “dry” states within a pore will depend on the balance of surface tensions between the liquid/vapor and liquid/protein interfaces. In particular, the water models used in many common biomolecular force fields (e.g. TIP3P, SPC/E, and TIP4P ^23^) significantly underestimate the liquid/vapor surface tension of pure water ^24^, which is a central parameter in this process ^8^.

The potential deficiency of the pairwise additive approximation is recognized, and over the last two decades a number of polarizable force fields have been developed for water as well as for biomolecular systems ^25^. Prominent amongst these are the AMOEBA force field, which uses polarizable atomic multipoles to simulate the effects of polarization within a classical MD framework ^26^, and the CHARMM Drude ^27^ force field, which employs massless Drude oscillators to model the electronic degrees of freedom of heavy atoms. The original AMOEBA03 water model reproduces many thermodynamic properties of water over a wide range of temperatures and pressures ^28–29^, and the more recent AMOEBA14 model ^30^ further improves upon this to reliably predict the properties of liquid- and gas-phase water for a large portion of its phase diagram. Both the AMOEBA ^31^ and the SWM4-NDP (Drude) ^32^ models also accurately describe the thermodynamics of ion solvation. Furthermore, the AMOEBA polarizable forcefield has recently been shown to yield a more accurate estimate of the electric field within the active site of an enzyme, as shown by correlation with quantum mechanically calculated electric fields compared to additive models ^33^.

Considering the potential of such polarizable force fields to more accurately predict the behavior of water, we therefore decided to quantify the effects of induced polarization experienced by water within an ion channel and its impact on the predicted hydration equilibrium of such pores. In this study, we now describe protocols for polarizable MD simulation of membrane protein (ion channel) systems and compare the pore hydration behavior of polarizable water models to that of fixed point-charge models. In addition to quantifying the dipole moment of water in the hydrophobic confinement of an ion channel pore, we also examine the influence of induced polarization on the energetics of ion conduction.

Our results demonstrate that although current non-polarizable approaches to the prediction of hydrophobic gates appear robust, there are cases where induced polarization may make more accurate predictions. A qualitative change in single-ion permeation energetics is observed when a polarized model is employed. Furthermore, changes in the solvation behavior of Cl^−^ ions within a hydrophobic channel are seen when polarizability is included in the model. We therefore discuss the significance of our findings in the context of ion channel function as well as their impact on the development of more accurate tools for the functional annotation of ion channels and other nanopores.

## Methods

### Equilibrium Molecular Dynamics Simulations

The majority of the simulations employed the pore-lining M2 helix bundle (residues 247 to 271) of the 5HT3R embedded in a dioleoylphosphatidylcholine (DOPC) bilayer. Additional simulations based on the 4PIR structure were conducted using either the M2 helix bundle in a palmitoyloleoylphosphatidylethanolamine (POPE) bilayer or the full TM domain (residues 220 to 334, and 426 to 459) in a DOPC bilayer. Equilibrium MD simulations were performed for four structures (i.e. PDB IDs 4PIR, 6BE1, 6DG7, and 6DG8), while umbrella sampling simulations of ion permeation were only carried out for the 6DG8 structure.

To assess the hydration behavior of different water models in the hydrophobic pore of the 5HT3R, a series of equilibrium MD simulations were carried out. Productive simulations using the additive CHARMM36m force field had a total duration of 150 ns, while for the more computationally intensive AMOEBA force field, the simulation time was 50 ns. Three independent repeats were carried out for each unique combination of parameters and the first 10 ns of each simulation were regarded as equilibration period and not included in the analysis. The details of the respective simulation protocols are described below.

### Structural Model and System Preparation

Ion channel structures were obtained from the PDB and missing atoms were added using the WHAT IF tool ^34^. The channel protein was then embedded in a homogeneous lipid bilayer using a serial multiscale procedure ^35^. Following a coarse-grained simulation to embed the protein in a lipid bilayer ^36^, the protein-bilayer system was converted back to an atomistic representation and re-solvated in a ~150 mM NaCl solution. This was followed by a 10 ns simulation with the additive CHARMM36m force field ^37^, the corresponding lipid parameters ^38–39^, and the CHARMM-modified mTIP3P water model ^40^ using the same parameters as the production simulations described in the following section.

### CHARMM Additive Force Field Simulations

As a basis for comparison, atomistic simulations using the additive CHARMM36m protein force field with associated lipid parameters and the mTIP3P water model were performed. Additionally, the CHARMM36m force field was combined with a number of further fixed point-charge water models. A comprehensive comparison of all existing additive water models is beyond the scope of this study. Instead, the SPC/E ^41^ and TIP4P ^23^ models were chosen as examples of widely used yet older models and the more recent TIP4P/2005 ^42^ and OPC ^43^ models were employed due to their improved accuracy in modelling water. The majority of the additive simulations focused on the mTIP3P and TIP4P/2005 models.

These simulations were carried out in GROMACS 2018 (http://www.gromacs.org/) using a leapfrog integrator with a step size of Δt = 2 fs for time integration. The length of bonds involving hydrogen atoms was constrained through the LINCS algorithm ^44^ and water geometry was kept rigid using the SETTLE ^45^ method. To preserve the experimentally determined protein structure, a harmonic restraining potential of 1000 kJ/mol/nm^2^ was applied to all Cα atoms.

The smooth PME method ^46^ with a real-space cutoff of 1 nm, a Fourier spacing of 0.12 nm, and charge interpolation through fourth-order B-splines was used to calculate electrostatic interactions under periodic boundary conditions. Van der Waals interactions were smoothly switched off between 1.0 and 1.2 nm and a long-range dispersion correction was applied to both energy and pressure. Simulations were performed in the isothermal-isobaric ensemble at a temperature of 310 K and a pressure of 1 bar. A velocity rescaling thermostat ^47^ with a coupling constant of τ_T_ = 0.1 ps and a semi-isotropic Parrinello-Rahman barostat ^48^ with a coupling constant of τ_P_ = 1 ps and a compressibility of 4.5 × 10^−5^ bar^−1^ were used for temperature and pressure control respectively.

### AMOEBA Force Field Simulations

Polarizable atomic multipole simulations were carried out in OpenMM 7.1.1 (http://openmm.org/) using the AMOEBA13 protein parameter set ^26^ in conjunction with both the AMOEBA03 ^28^ and AMOEBA14 ^30^ water models, as well as polarizable multipole parameters for ions ^31^ and lipids ^49^. Starting configurations for these simulations were derived from the final frames of a 10 ns long atomistic equilibration simulation as described above. Production simulations were preceded by 1000 steps of energy minimization with the AMOEBA force field to avoid divergent energies due to induced dipoles.

Time integration was performed using the r-RESPA method ^50^ with an outer time step of Δt = 2 fs and an inner time step of δt = 0.25 fs. Electrostatic multipole and van der Waals forces were updated in intervals of the outer time step; all other forces were evaluated at each inner time step. OpenMM was operated in mixed-precision mode with forces being accumulated in single precision and time integration performed in double precision.

Electrostatic multipole interactions were treated through the PME method ^51^, with a real-space cutoff distance of 0.8 nm, an Ewald error tolerance of 5 × 10^−4^, and interpolation performed through fifth-order B-splines. Van der Waals forces were computed up to a distance of 1.2 nm and a long-range dispersion correction was applied to account for interactions beyond this cutoff ^52^. Induced dipoles were computed through truncated self-consistency iteration using the extrapolated perturbation theory method ^53^.

The simulation system was maintained at a temperature of 310 K using an Andersen thermostat with a collision frequency of 1 ps^−1^. An anisotropic Monte Carlo barostat was used to enforce a system pressure of 1 bar with Monte Carlo moves attempted every 20^th^ integration step. In accordance with the AMOEBA model, no constraints on covalent bonds or water molecule geometry were enforced. However, in order to restrain the protein in a configuration close to its experimentally determined structure, all Cα atoms were placed under a harmonic restraint with a force constant of 1000 kJ/mol/nm^2^.

### Umbrella Sampling

Umbrella sampling was performed to obtain one-dimensional single-ion potential of mean force profiles for both Na^+^ and Cl^−^ ions passing through the pore of the open-state structure (PDB ID: 6DG8) of the 5HT3R. These simulations were carried out using the AMOEBA force field with the AMOEBA14 water model, or the CHARMM36m force field with either the mTIP3P or TIP4P/2005 water models. Simulation protocols were similar to those detailed above for equilibrium MD simulations.

The collective variable was chosen to be the distance between the ion and the center of mass of the protein along the *z*-axis of the simulation box, which corresponds to the direction normal to the lipid bilayer membrane. Starting configurations for the umbrella windows were generated from the final state of the equilibrium MD simulations. A target ion was relocated to subsequent positions along the *z*-axis and the lateral position of the ion was set to the centre of mass of the protein to ensure a placement inside the channel pore. This was followed by 10 steps of energy minimization to remove steric clashes between the target ion and water molecules present in the pore. During both minimization and sampling, a harmonic biasing force of 2000 kJ/mol/nm^2^ was applied to restrain the collective variable.

Umbrella windows covered the entire length of the ion channel and reached up to 1 nm into the bulk water regime. Each umbrella window was simulated for 10 ns. The distance between two subsequent umbrella windows was 0.1 nm. This setup corresponds to 62 windows with a cumulative simulation time of 620 ns for each of Na^+^ and Cl^−^. Unbiasing was performed through the weighted histogram analysis method (WHAM) using the Grossfield lab implementation in version 2.0.9 (http://membrane.urmc.rochester.edu/wordpress/?page_id=126). The first half of each simulation window was regarded as equilibration period and the final PMF profile, *F(z)*, was calculated only from the final 5 ns of simulation time.

## Results and Discussion

### Simulation system

As a model system, we chose to examine the behavior of polarizable water within the transmembrane pore of the 5-HT_3_ receptor (5HT3R), a pLGIC that opens upon binding of serotonin (5-HT_3_) (**Figure 1**). The pore of this channel has previously been shown to contain a hydrophobic gate at the 9′ position in the pore-lining M2 helix ^15–16, 54^. In previous studies we have shown that a reduced protein system focused on these pore-lining M2 helices of the 5HT3R channel, embedded in a phospholipid bilayer, provided an accurate model system; we therefore chose a similar approach. This reduced system (**Figure 1A**) also allows several different conformational states of the channel to be readily compared, e.g. closed, intermediate, and open states (see below and ^16^). Importantly, simulations in the presence of an applied electrostatic field were used to compare ion fluxes through the pentameric bundle of M2 helices used here with a more complete protein model that includes the entire transmembrane domain (TMD) of the 5HT3R. The two systems generated a similar level of conductance (corresponding to ~30 pS), with selectivity of sodium over chloride ions (see **SI Fig. S1**), thus supporting our rationale for using this reduced M2 helix bundle in the current studies.

**Figure 1:**
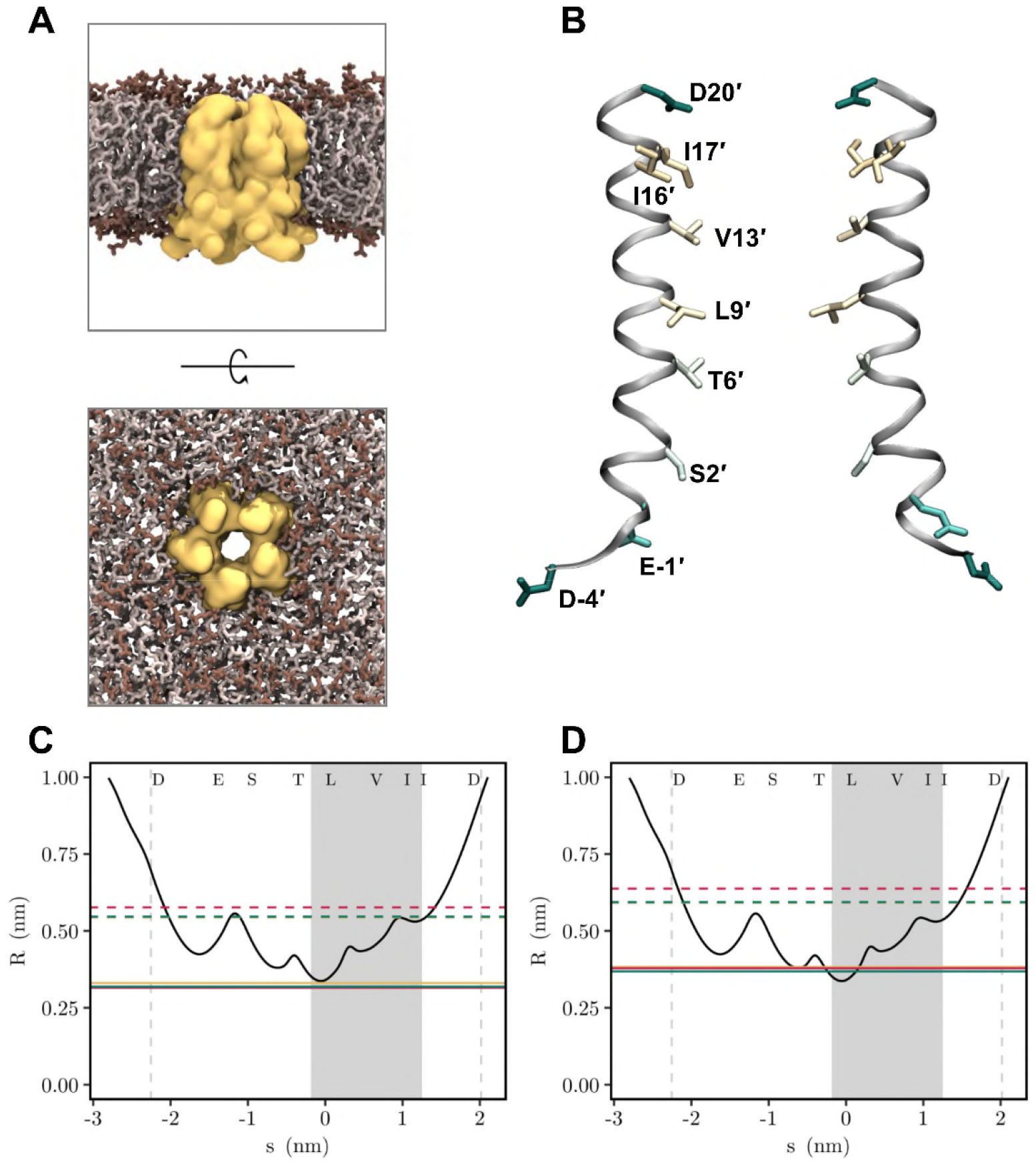
**A** Protein position relative to lipid bilayer after equilibration with the AMOEBA polarizable multipole force field. Shown is a surface representation of the M2 helix bundle of the 5HT3R structure (PDB ID 6DG8; yellow) embedded in a DOPC bilayer as viewed from the side and from the extracellular mouth of the TM domain. The protein-bilayer system is shown at the end of a 10 ns simulation using the AMOEBA polarizable multipole force field which was in turn initiated using coordinates resulting from equilibration with the CHARMM36m force field. Lipids are displayed in liquorice representation with headgroups and hydrocarbon tails colored in dark and light shades of brown respectively. Water molecules, ions, and lipid hydrogens are omitted for clarity and lipid molecules obstructing the view of the protein are not shown. **B** The M2 helix bundle in the open-state (PDB ID: 6DG8) structure of the 5HT3R receptor. Two of the M2 helices are shown as ribbons with pore-facing residues displayed in liquorice representation and colored by hydrophobicity. **C, D** Pore radius profile (black curved line) of the open-state (PDB ID: 6DG8) structure of the 5HT3R compared with the radii of the first (solid lines; yellow = AMOEBA14; red = mTIP3P; green = TIP4P/2005) and second (dashed lines) hydration shells of (**C**) Na^+^ ions and (**D**) Cl^−^ ions. Hydration shell radii were calculated from the minima of the radial distribution functions (see **SI Fig. S6**). The shaded area and vertical dashed lines represent the hydrophobic gate region and the extent of the protein respectively.

### Influence of polarization on the behavior of water in a hydrophobic gate

Having validated our system, we next compared pore hydration predicted by the AMOEBA polarizable MD simulations with the additive CHARMM36m force field combined with either the mTIP3P or TIP4P/2005 water model. Consistent with previous studies ^15, 54 16^, the additive MD simulations predict complete de-wetting of the hydrophobic gate in both the closed (PDB ID: 4PIR and 6BE1) and intermediate (PDB ID: 6DG7) state structures. Both polarizable water models were in good agreement with this (**Figure 2A**). Furthermore, in the remainder of the pore, there was also a high level of agreement between the different water models which all predicted a water density close to that of bulk water. The time evolution of the phase state of water within the hydrophobic gate region is shown in **Figure 2B**. This reveals that in the closed and intermediate state structures, water remains in a vapor state throughout the simulation, with only a few brief wetting events occurring in some repeats. This was consistent across all four water models, indicating that the free energy cost of hydrating these particular pore conformations is particularly high.

**Figure 2:**
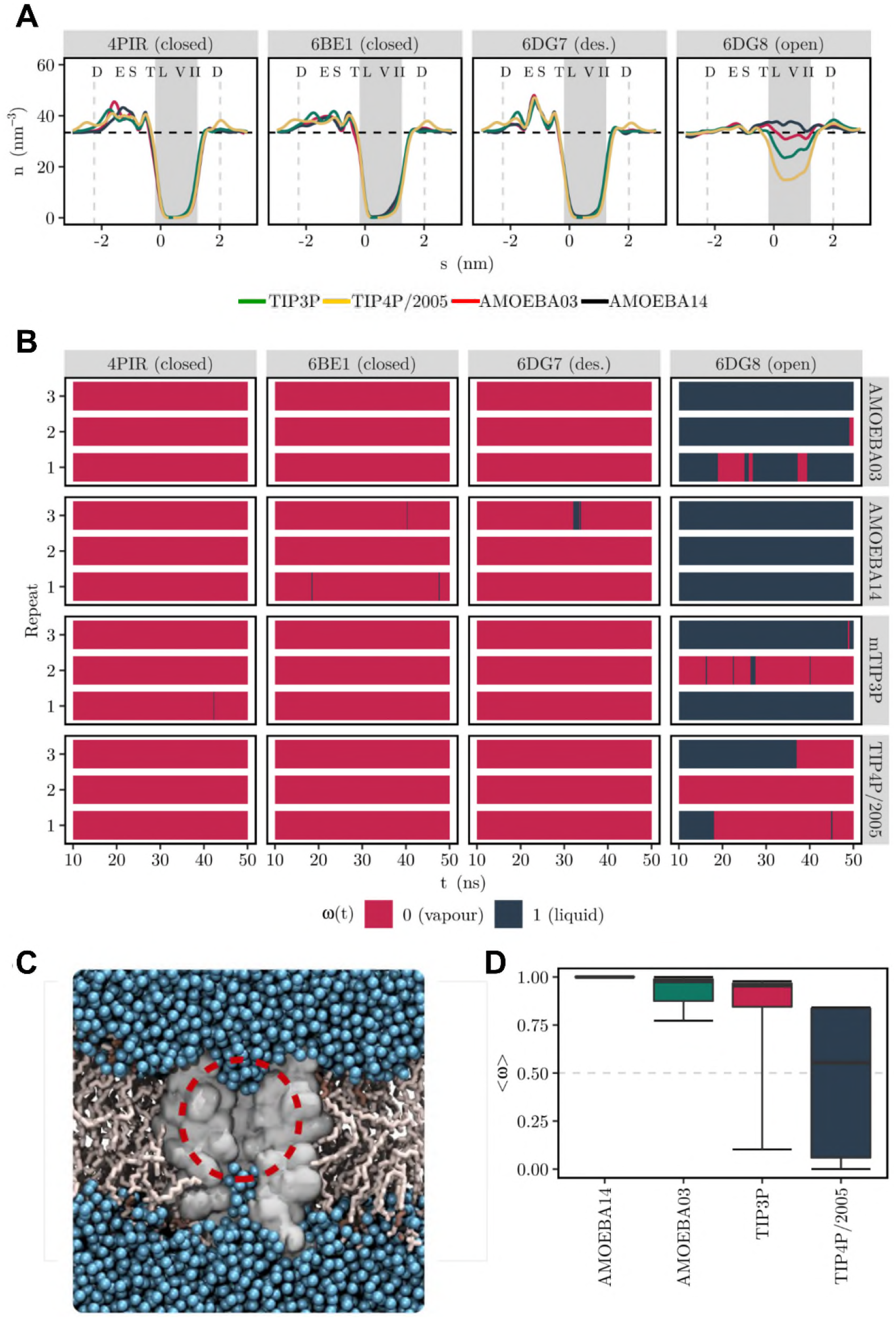
Comparison of pore hydration using additive and polarizable force fields and water models. **A** Water density in four conformational states of the 5HT3R: PDB IDs 4PIR and 6BE1 are closed states, 6DG7 is an intermediate state and 6DG8 is an open state. Profiles are shown for mTIP3P (green), TIP4P/2005 (yellow), AMOEBA03 (red) and AMOEBA14 (black). The profiles shown represent the average of three independent simulations (five in the case of mTIP3P) and the dashed horizontal line indicates the density of bulk water. Dashed vertical lines and the shaded background denote the extent of the protein and the hydrophobic gate region respectively. Single-letter codes for the pore-facing amino acid side chains are given at the top of each panel. **B** Beckstein openness over time for three independent repeats. Repeats 4 and 5 are not shown for the mTIP3P water model and the final 100 ns of the mTIP3P simulations are omitted for clarity. **C** Snapshots of the de-wetted pore from simulation of the closed state 5HT3R pore (PDB ID 4PIR) using the mTIP3P water model. The protein and lipid molecules are in grey; the waters in blue. The red circle highlighted the de-wetted region of the pore. **D** Hydration equilibrium of the open-state (PDB ID: 6DG8) 5HT3R pore predicted by the different water models. The time-averaged Beckstein openness, <ω>, is shown as a measure for the fraction of simulation time during which the channel pore is hydrated. The boxplot shows the median and quartiles across five independent repeats for the additive and three independent repeats for the polarizable water models.

However, when we examined the behavior of water in a recently solved structure of 5HT3R (PDB ID: 6DG8) that was predicted to be open by computational electrophysiology (see **SI Fig. S1**) ^16^, we observed marked differences between these water models. The additive simulation predicted significant de-wetting within this hydrophobic pore region (**Figure 2A**). Although with a time-averaged water density of ~50% of bulk water, this region remained substantially more hydrated than the closed and intermediate (6DG7) states of the receptor. Both additive water models also produced stochastic fluctuations between the liquid and vapor states within the hydrophobic gate region (**Figure 2B**). By marked contrast, we found that during simulations of the 6DG8 structure with the AMOEBA polarizable force field, water strongly favored the liquid (i.e. wetted) state, with the time-averaged water density being close to that of bulk water throughout the entire channel pore (**Figure 2A** **and** **2B**). In all three simulations with the AMOEBA14 water model, the hydrophobic gate remained hydrated with its waters exclusively in the liquid state. It is therefore expected that such a fully hydrated pore does indeed represent an open, i.e. conductive, state of the channel.

We also investigated whether alternative additive water models exhibit a qualitatively similar behavior. As shown in **SI Fig. S3**, we examined the pore hydration behavior in simulations employing the OPC, TIP4P, TIP4P/2005, and SPC/E water models in conjunction with the CHARMM36m force field. Simulations of the closed (4PIR and 6BE1) and intermediate (6DG7) state structures mirrored the behavior observed in polarizable MD simulations. For the open state (6DG8) pore liquid-vapor transitions were observed with all these additive water models. The time-averaged water density in the hydrophobic gate region varied between zero and the density of bulk water.

It is therefore evident that the only unambiguous prediction of a fully hydrated pore in the 6DG8 structure is made by using what is possibly the most accurate water model employed. With the exception of AMOEBA14, all water models undergo liquid-vapor transitions in the pore of the 6DG8 structure. Consequently, comparison of simulations using this particular 5HT3R conformation can be used to establish a ranking of water models with respect to their relative proclivity for the formation of liquid or vapor states. **Figure 2D** displays the time-averaged openness for each water model for the open state of the channel. This reveals that the two polarizable AMOEBA models strongly favor the liquid state, whilst amongst the additive models, mTIP3P is the most likely to predict a hydrated pore.

### Polarization effects on nano-confined water

Having established that induced polarization can influence predictions of how water behaves in the nano-confined environment of an ion channel pore, we next attempted to quantify these effects in more detail. **Figure 3A** shows how the magnitude and orientation of the molecular dipole moment of water vary along the pore of the 5HT3R. As expected, the additive models do not exhibit variation in any of the pore structures examined. Likewise, when examined with the AMOEBA models, the closed (4PIR and 6BE1) and intermediate (6DG7) state structures all yielded very similar profiles across the TMD region. In these three structures, both AMOEBA models exhibit a decreased dipole moment within the hydrophobic gate region. Here the dipole moment drops from ~2.7 D in the bulk to a minimum of ~2.2 D between the 9′ leucine and 13′ valine residues where de-wetting typically occurs. Consistent with the idea that water adopts the vapor phase within this region, the dipole moment of water is effectively reduced to a value close to that predicted for its gas phase.

**Figure 3:**
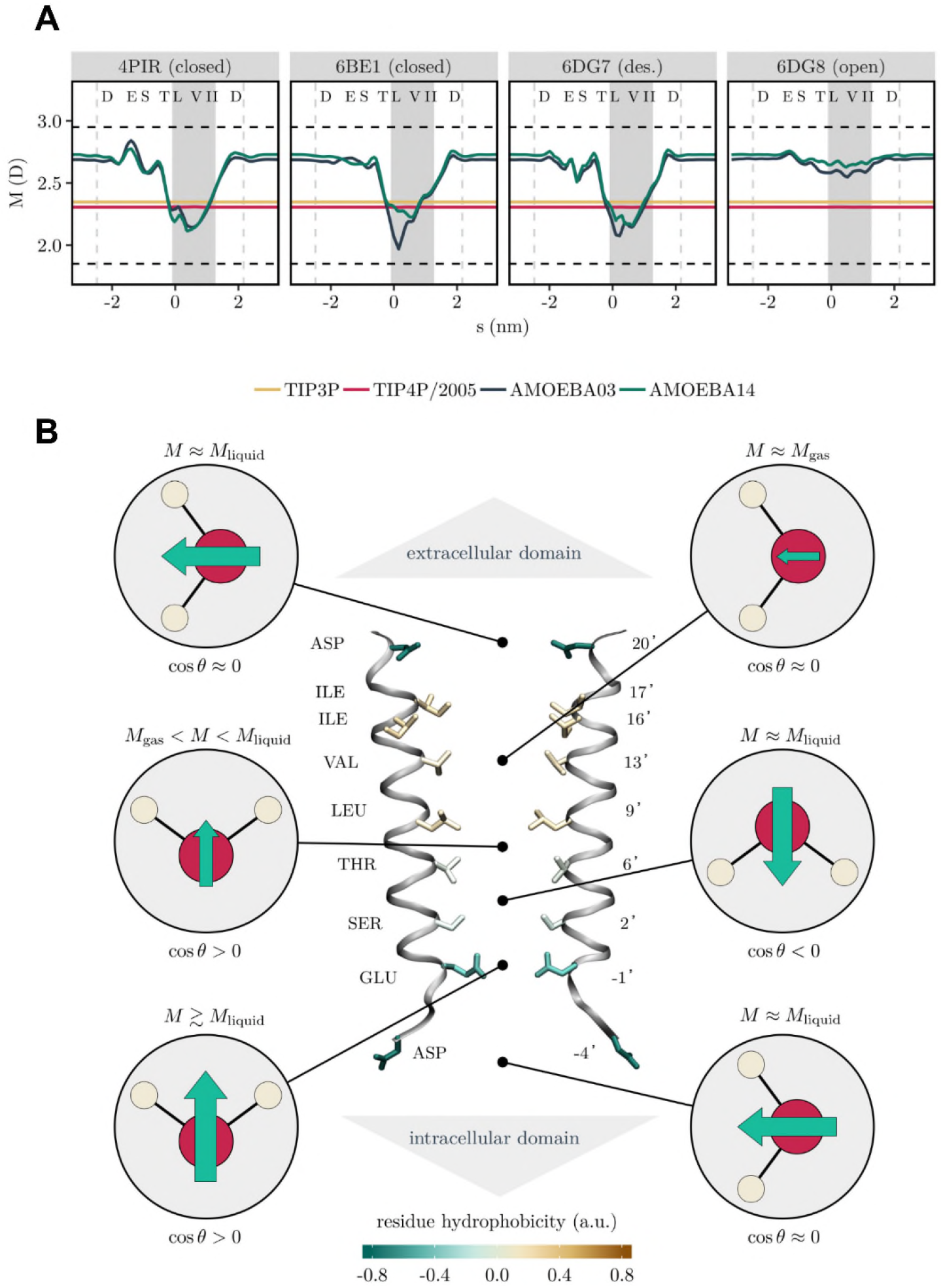
**A** Magnitude of the molecular dipole moment, *M*, of water in the pore of four conformational states of the 5HT3R. The shaded area represents the hydrophobic gate region, in which de-wetting occurs. Dashed vertical lines indicate the extent of the simulated protein structure. Dashed horizontal lines at 1.85D and 2.95D represent the dipole moment of water in the gas and liquid phase respectively. Single-letter codes for the pore-facing amino acid side chains are given at the top of each panel. Curves shown represent the average over three independent repeats. **B** Schematic summary of water dipole moment behavior in the closed-state (PDB ID: 4PIR) structure of the 5HT3R. Two of the M2 helices are shown as ribbons with pore-facing residues displayed in liquorice representation and colored by hydrophobicity. At six positions along the channel pore, the characteristic magnitude and orientation of the molecular dipole moment of water (for the AMOEBA models) are indicated by a green arrow representing the net dipole vector. In the hydrophobic pore section between the 9′ leucine and 16′ isoleucine residue rings, the dipole moment is significantly reduced relative to its value in bulk water. Here the water dipole resembles that of gas-phase water molecules.

Interestingly, the effect of the hydrophobic gate on the dipole moment is much larger than that of charged residues lining the pore. For example, in the closed state structure (4PIR), the dipole moment of both AMOEBA models peaks near the −1′ glutamate residue (**Figure 3A**), but this peak is only a fraction of the decrease experienced within the hydrophobic gate. Furthermore, in the other two de-wetted structures (6BE1 and 6DG7), this peak is absent, but the large reduction in dipole moment within the 9′ hydrophobic gate region remains. This emphasizes the importance of the hydrophobic nature of the 9′ leucine and nearby hydrophobic residues as the principal gate within the 5HT3R pore. Consistent with this, and with our prediction that the 6DG8 structure represents a fully open conformation, the decrease in dipole moment throughout the hydrophobic section of this pore is relatively small, with both AMOEBA models exhibiting only small fluctuations around the bulk value. Even the intermittent de-wetting events occurring during the third AMOEBA03 repeat (**Figure 2B**) are also reflected here by a slightly lower dipole moment.

The nearly constant dipole moment profile in the open-state 6DG8 structure also suggests that the sharp decrease in dipole moment in the closed- and intermediate-state structures is not simply caused by the presence of hydrophobic amino acids, but instead due to the absence of neighboring water molecules. The hydrophobicity of the pore surface is very similar in all these structures of 5HT3R, but in the open-state structure the dipole moment of water decreases by no more than ~0.1 D even in direct proximity to a ring of leucine residues. Furthermore, the orientational behavior of the dipole moment is consistent across both polarizable models and is similarly captured by the additive mTIP3P model (**SI Fig. S4A**).

To explore whether these dipole profiles are sensitive to the overall electrostatic environment experienced by the protein, two additional sets of 5HT3R simulations were conducted based on the closed-state structure (4PIR), where we varied the charge composition of the lipid bilayer and/or protein. In the first, the M2 helix bundle was embedded in a POPE rather than a DOPC lipid bilayer, whilst in the second, the complete transmembrane (TM) domain of the channel was embedded in a DOPC bilayer. Throughout both sets of simulations the hydrophobic gate remained de-wetted, and comparison of the corresponding dipole moment profiles revealed all the profiles derived using the AMOEBA force field were in near-quantitative agreement (**SI Fig. S4B**). This confirms that the dipole moment of water inside the pore is primarily influenced by short-ranged interactions. These effects are summarized in **Figure 3B**.

Upon entering the pore from the intracellular side (i.e. *s* < −1.5 nm) the dipole moment is enhanced near the −1′ ring of glutamate (E) residues. Further into the channel (around *s =* −0.5 nm) its magnitude returns to the bulk value and its orientation is flipped. As water approaches the 9′ ring of leucine residues (around *s* = 0 nm), the magnitude of the dipole moment decreases markedly and its orientation reverses again. In the middle of the hydrophobic gate region (around *s* = +0.5 nm) the dipole moment approaches a value similar to that of gas-phase water and loses its orientational preference.

### Influence of polarization on the energetics of ion permeation

We next examined how the behavior of ions is impacted by induced polarization and what role these different water models play in the energetics of ion conduction. To determine this, umbrella sampling simulations were carried out using the open-state (6DG8) structure to obtain a potential of mean force (PMF) profile for both single Na^+^ and Cl^−^ ions (**Figure 4**). These simulations employ the polarizable AMOEBA force field with the AMOEBA14 water model, compared with the additive CHARMM36m force field with the mTIP3P and the TIP4P/2005 water models. The additive mTIP3P and TIP4P/2005 water models yield qualitatively similar profiles for both ions. Examination of the umbrella histograms also reveals good overlap and error metrics, indicating that the resulting PMF profiles are converged (**SI Fig. S5**).

**Figure 4:**
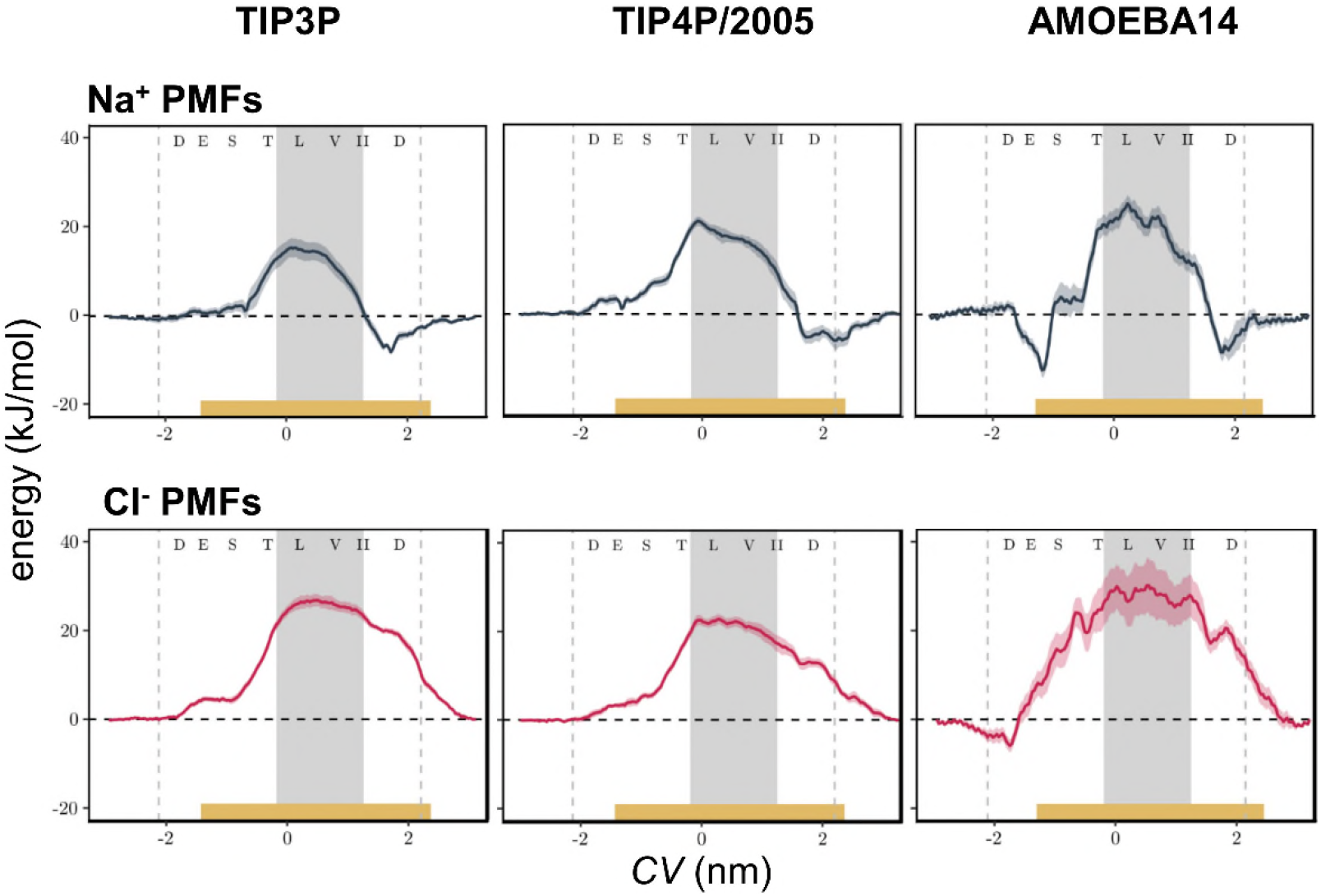
Single-ion PMF profiles for Cl^−^ (red) or Na^+^ (black) ions in the pore of the open-state 5HT3R (PDB ID: 6DG8). The solid line indicates the free energy profile calculated from the final 5 ns of each umbrella window. Confidence bands were obtained by calculating the standard error over independent 1 ns sampling blocks during the same time period. Vertical dashed lines denote the extent of the protein and the yellow bar represents the position of the lipid bilayer. The hydrophobic gate region, where de-wetting occurs in equilibrium simulations, is denoted by the grey background shading. Single-letter codes for the pore-facing amino acid side chains are given at the top of each panel. Dashed horizontal lines indicate the bulk energy. The collective variable (CV) is defined as the distance along the z-axis between the ion and the protein center of mass and thus is zero at the center of the pore.

For Na^+^, the additive PMFs are characterized by a free energy well near the 20′ aspartate residues and a barrier within the hydrophobic gate region, but there is no energy well near the −1′ ring of glutamate residues. For both additive water models, the energy barrier also extends from the 6′ threonine to the 16′ isoleucine residues. Quantitatively, the energy barrier for the TIP4P/2005 model (21.0 ± 1.1 kJ/mol), is higher than the value for the mTIP3P model (15.6 ± 2.2 kJ/mol). However, Cl^−^ experiences the largest energy barriers within the hydrophobic gate region (26.9 ± 1.4 kJ/mol and 22.8 ± 1.1 kJ/mol with the mTIP3P and TIP4P/2005 models respectively). Both water models thus predict a larger barrier for Cl^−^ than Na^+^, which correlates with the experimental cation selectivity of the 5HT3R ^55^.

Interestingly, free energy profiles derived using AMOEBA exhibited notable differences from the additive models (**Figure 4**). For Na^+^, the profile still peaked within the hydrophobic gate (25.0 ± 1.9 kJ/mol near the 9′ leucine). However, it also revealed two energy wells near the 20′ aspartate and −1′ glutamate residues. The profiles also appeared more rugged and exhibited more secondary features on a smaller spatial scale. This suggests that for a polarizable ion there may be preferential interactions along the channel pore, depending on whether it can create induced dipoles in neighboring atoms.

Overall, the Cl^−^ PMF derived from the AMOEBA force field resembles that predicted by the mTIP3P model and peaks in the hydrophobic gate region (30.2 ± 6.2 kJ/mol compared with 26.9 ± 1.4 kJ/mol for mTIP3P). However, AMOEBA predicts a more gradual increase of the PMF and a small energy well in the vicinity of the −4′ ring of aspartate residues. This is likely due to the fact that these charged side chains do not point directly towards the pore, so Cl^−^ can favorably interact with the backbone atoms, in which it may induce additional polarization. Convergence of the AMOEBA PMFs also required more time (~5 ns for Na^+^ and ~6 ns for Cl). However, beyond the need for longer equilibration periods, these effects may also be relevant for the dynamics and selectivity of ion permeation as the differential interactions of polarizable ions with the pore lining may not only alter the rate of transport but also its anion vs. cation selectivity. Specifically, a polarizable ion that interacted with the pore walls more strongly would experience a larger effective friction and would consequently be transported through the channel at a lower rate than a non-polarizable ion, provided that the driving force was identical.

### Ion hydration within the pore

In order to better understand the interaction of polarizable ions with the pore lining, it is also important to consider changes in ion hydration during movement through the pore. We therefore calculated radial distribution functions (RDFs) of water oxygens around Na^+^ ions and Cl^−^ ions in bulk solution (**SI Fig. S6**). For all of the water models around Na^+^ the RDFs exhibit a sharp first peak at ~0.25 nm and a second broader peak at ~0.45 nm. These two peaks are clearly separated and the RDF approaches zero after the first peak, thus defining the first hydration shell. The AMOEBA14 model predicts a first hydration shell for Na^+^ of 6.0 waters, whilst the mTIP3P and TIP4P/2005 models predict of 5.8 and 5.9 respectively. A hydration number of 6.0 has also previously been reported for the AMOEBA03 model ^31^. By comparison, predictions from recent *ab initio* MD simulations fall in the range from 5.1 to 6.1 ^56–57^. Older experiments ^58^ predict the first hydration shell of Na^+^ ions to range from 4.0 to 8.0, whilst more recent LAXS experiments yield a value of 6.0 ^59^. As anticipated, Cl^−^ has an RDF (**SI Fig. S6**) that is shifted towards greater separations than the corresponding sodium-oxygen RDFs. All three water models yield similar values for the location of the first RDF peak and the radius of the first hydration shell. The first-shell hydration number for Cl^−^ is predicted to be 6.6 for both the AMOEBA14 and TIP4P/2005 models, and 7.4 for the mTIP3P model. These exceed the previously reported value of 6.0 for the AMOEBA03 water model ^31^. Both the AMOEBA14 and TIP4P/2005 models fall within the 6.3-6.6 range predicted by *ab initio* calculations ^57^. Neutron scattering predicts a hydration number of 7.0 ± 0.4 ^60^ whilst earlier experiments indicate values between 5.3 and 6.2 ^61^. Thus, overall, all three water models yield first-shell hydration numbers consistent with other available data.

To estimate how these hydration shells change during ion permeation, we compared them to the pore radius profile for the open (6DG8) channel structure (**Figure 1CD**). The first hydration shell of Na^+^ is smaller than the narrowest constriction of the open 6DG8 structure for all 3 water models (**Figure 1C**). However, the radius of the second hydration shell exceeds the pore radius nearly everywhere. Consequently, Na^+^ can remain partially hydrated as it traverses the pore, but is expected to lose part or all of its second hydration shell. Irrespective of which water model is used, even the first hydration shell of Cl^−^ is slightly larger than the narrowest constriction at the 9′ leucine position. However, the first hydration shell should remain intact throughout the remainder of the pore. Similar to Na^+^, the second hydration shell of Cl^−^ cannot be accommodated within the pore, regardless of the water model used.

The hydration number profiles for Na^+^ ions within the open (6DG8) state 5HT3R pore (**Figure 5AB**) show that for each of the three water models, the first shell hydration number decreases by less than ~0.5 (compared to their bulk values) when the ion is within the hydrophobic gate. This is consistent with an otherwise intact inner hydration shell. However, for the AMOEBA model this value is markedly reduced near the −1′ glutamate and 20′ aspartate residues. Similar but smaller decreases were also observed for the mTIP3P and TIP4P/2005 models. Snapshots of the simulation reveal this can be explained by interactions with one of the negatively charged side chains which attract the ion into a position close to the pore surface. This in turn displaces a significant fraction of the first hydration shell (**Figure 5A**). While this effect also appears to exist for the additive mTIP3P and TIP4P/2005 models, it is significantly enhanced for the polarizable AMOEBA force field.

**Figure 5:**
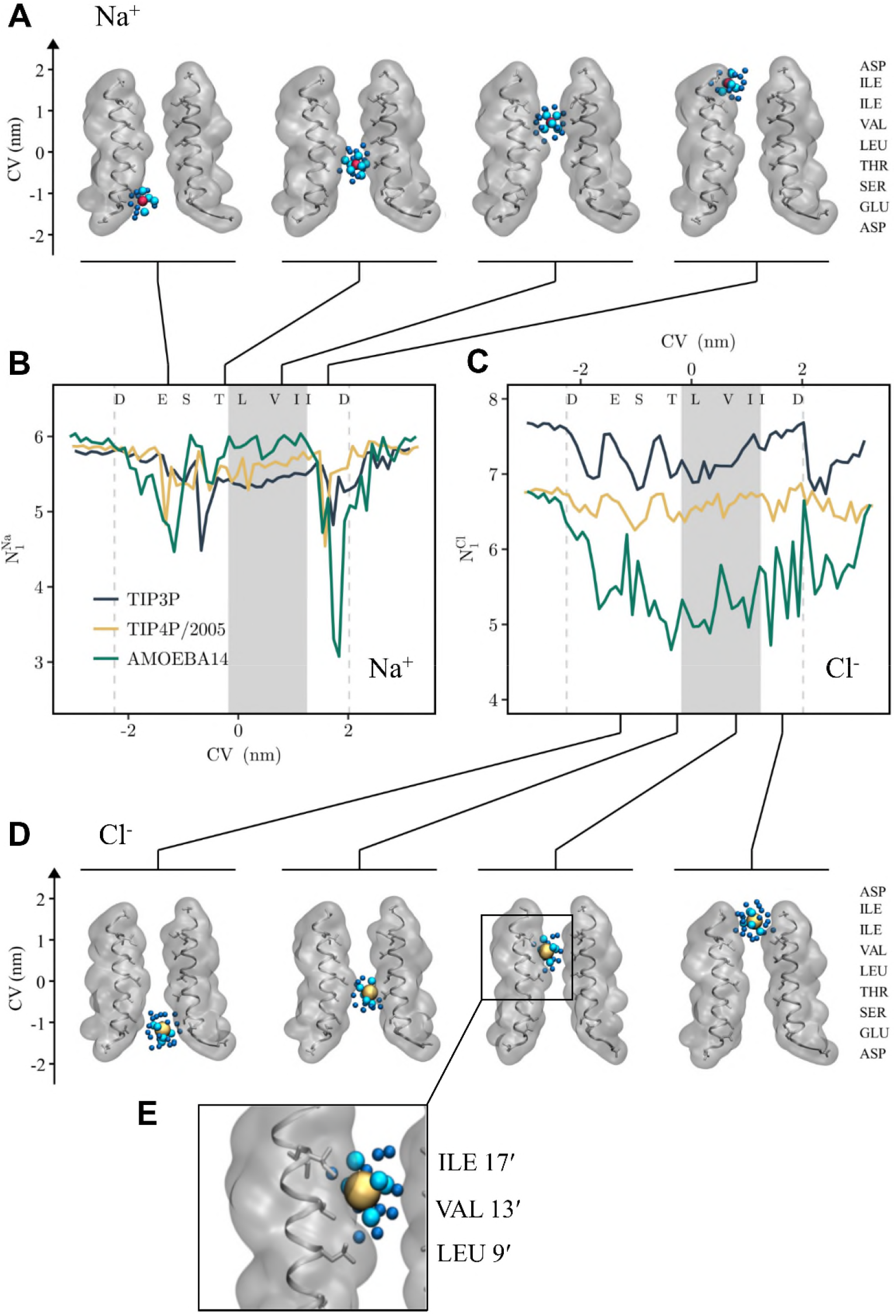
Hydration number profile for (**A,B**) Na^+^ ions and (**CDE**) Cl^−^ ions. **B** Variation of the first-shell hydration number of Na^+^ ions along the pore of the open-conformation 5HT3R (PDB ID: 6DG8). The shaded area and vertical dashed lines represent the hydrophobic gate region and the extent of the protein respectively. The position of the snapshots shown in **A** is indicated by the linking black lines. Representative snapshots are shown (**A**) from four umbrella sampling windows employing the AMOEBA14 force field. The restrained Na^+^ ion is shown as a red van der Waals sphere, while oxygens of water molecules in the first and second hydration shell are shown in cyan and blue respectively. Two of the five pore-lining protein helices are shown in both cartoon and surface representation. **C,D** Variation of the first-shell hydration number of Cl^−^ ions along the pore of the open-conformation 5HT3R. Representative snapshots are shown (**D**) from four umbrella sampling windows employing the AMOEBA14 force field. The restrained Cl^−^ ion is shown as a yellow van der Waals sphere, while oxygens of water molecules in the first and second hydration shell are shown in cyan and blue respectively. **E** is a zoomed-in image of a Cl^−^ ion interacting with three hydrophobic side chains (see main text for details).

By marked contrast, the hydration profiles for Cl^−^ (**Figure 5C**) reveal a clear difference between water models. With both additive models, the shell remains mostly constant along the entire length of the pore and only exhibits local fluctuations of ~0.5. However, the polarizable model predicts a marked decrease throughout the pore, falling from a bulk value of ~6.5 to ~5.0 within the hydrophobic region of the pore. This behavior suggests that when a polarizable force field is used, Cl^−^ spends a longer time than Na^+^ close to the hydrophobic surface of the pore.

Polarizable Na^+^ and Cl^−^ ions therefore appear to exhibit a very different behavior in the vicinity of hydrophobic pore-lining side chains of the gate region of the 5HT3R pore. While Cl^−^ relinquishes part of its hydration shell to form a tight association with the hydrophobic surface, Na^+^ ions retain most of their hydration shell. This effect is not reproduced by additive force fields. The interaction of Cl^−^ ions with hydrophobic residues can be seen at a number of locations along the pore (**Figure 5D**) and at one position Cl^−^ ion is hydrated by 4 to 5 inner-shell waters on one side, but on the other side ~2 waters have been displaced and instead the ion interacts directly with a hydrophobic surface formed by the side chains of residues L9′, V13′ and I17′ (**Figure 5E**)

Hydrophobic interactions with Cl^−^ have been reported before. For example, high-resolution structures of a number of Cl^−^ transporters and channels have revealed Cl^−^ binding sites formed by a mixture of charged and hydrophobic contacts with Cl^−^. Well-defined examples of this are seen in the structures of e.g. halo rhodopsin (1E12) ^62^, the NTQ Cl^−^ transport rhodopsin (5G28) ^63^, and the bestrophin-1 chloride channel (4RDQ) ^64^. Furthermore, a number of simulation studies of ions at water/air interfaces have shown that Cl^−^ (and Br^−^) are found at the water/air interface, unlike Na^+^ which remains fully hydrated and so avoids the interface ^65–67^. Interestingly however, this anion accumulation at the water/air interface is only seen if a polarizable model (or its equivalent, see e.g. ^20^) is employed. This is explained in terms of polarizable anion dielectric continuum theory (PA-DCT) where polarization of large anions (including Cl^−^) at this interface compensates for loss of water/ion H-bonds ^68^.

## Conclusions

A number of additive and polarizable water models have been comprehensively explored in terms of two main approaches to the functional prediction of ion channel pore properties, namely hydrophobic gating and ion permeation free energy profiles. The results provide important insights into how polarizable models may allow us to alter and/or refine our understanding of the physicochemical and structural basis of ion channel function.

The comparison of the three different water models for four different conformational states of the 5HT3R channel demonstrates that previous conclusions derived about hydrophobic gating in these channels are quite robust regarding the choice of water model. This is important as such simulations are now being increasingly employed to functionally annotate the rapidly increasing number of ion channel structures that are being solved ^6, 15, 18^. Our study indicates that such simulation-based annotation of potential hydrophobic gates is qualitatively robust in distinguishing between closed and open states. However, for an open-state structure such as the 6DG8 conformation of 5HT3R which appears to be ‘on the edge’ of wettability there are quantitative differences between these water models, with AMOEBA14 most consistently predicting full solvation of the open state of the pore. In such cases where the free energy difference between liquid and vapor states is close to zero, it is perhaps not surprising that exact quantitative predictions vary considerably with the water model employed and require not only a water model whose bulk properties are in quantitative agreement with experiment, but also a correct description of surface tension at the water-protein interface.

As might be anticipated our simulations of ion permeation were also sensitive to the water models and force field employed. Whilst there was qualitative agreement between single-ion permeation free energy profiles calculated for the additive and polarizable models, there were also key differences. In particular, the polarizable models result in more rugged (i.e. detailed) PMF profiles, indicating that induced polarization results in more specific interactions between the ion and protein. In particular, our investigation of ion hydration suggests that with a polarizable force field, ions may interact with pore-lining residues more favorably, to produce a higher degree of friction and a lower predicted conductivity. Future studies might therefore address this by e.g. computational electrophysiology ^69^ simulations.

Umbrella sampling simulations employing a polarizable force field have so far only been reported for a small number of ion channel systems ^70^. A study of the gramicidin A channel ^71^ suggests that the explicit inclusion of polarization effects significantly improved computational predictions of ion channel conductance.

Free energy profiles for ions in the gramicidin A channel were calculated with the AMOEBA force field, suggesting that polarization effects reduced the energetic barrier experienced by ions relative to simulations with the CHARMM27 force field ^71^. However, gramicidin A forms a narrow (single-file) channel pore where the ion interacts mainly with protein backbone atoms (as is also the case in the selectivity filter of K^+^ channels ^70^). In the current study, we are examining channels in which ions permeate in a largely hydrated state and so are more likely to be sensitive to the water models employed. Such factors may therefore be particularly important for studies of Na^+^ and Ca^2+^ channels, where these ions are also thought to permeate in their fully or partially hydrated forms.

Whilst exploring permeation free energy profiles with AMOEBA we observed apparently favorable hydrophobic contacts of the protein with Cl^−^. This correlates with other recent studies of anions at air/water interfaces, with structural studies of Cl^−^ binding sites in Cl^−^ channels and Cl^−^ transport proteins, and with the properties of novel anionophores (biotin[6]uril hexaesters) which exploit C-H hydrogen bond donors in order to favor the transport of softer, more polarizable anions such as chloride over hard anions such as bicarbonate ^72^. This merits further investigations into how the use of polarizable force fields may modify our understanding of anion selectivity in biological ion channels.

Overall, our studies suggest that more extensive simulations using polarizable force fields on a wider range of channel structures, including direct simulations of ion permeation (e.g. computational electrophysiology) will provide new insights into the mechanisms of rapid ion permeation and high (anion) selectivity. This is a realistic aspiration given the relative scale of the simulations described above, as the computational cost of the AMOEBA simulations is only about one order of magnitude greater than for equivalent additive simulations.

## Acknowledgements

Our thanks for discussions with our colleagues concerning this work, especially Dr. Charlotte Lynch. This work was funded by grants from BBSRC, EPSRC, and Wellcome.

## Conflict of Interest

The authors declare that they have no conflict of interest.

