## Supplementary material for "Induced Polarization in MD Simulations of the 5HT_3_ Receptor Channel": SI Text and Figures

### Supplementary Methodological Detail

#### *Simulation protocols*

The parameters chosen for simulations with the CHARMM and AMOEBA force fields were selected to reflect the conditions under which the respective force field was originally parameterised. However, because relatively few studies have employed polarisable force fields to investigate membrane proteins, it was important to ascertain that these parameters permit stable integration and sampling from the appropriate thermodynamic ensemble. Thus, exploratory simulations were performed using the M2 helix bundle from the closed-state structure (PDB ID: 4PIR) of the 5HT3R embedded in a DOPC lipid bilayer as a test system. The total energy, box volume, and average temperature over the initial 0.25 ns of a 1 ns simulation were analysed for the AMOEBA forcefield (**SI Fig. S2**) for their stability. Additionally, the protein position relative to lipid bilayer after equilibration at the end of 10 ns long simulations using the additive CHARMM36 force field or the AMOEBA polarisable multipole force field showed similar positions of the M2 bundle in the bilayer. Together these results suggest that the system consisting of a pentameric M2 bundle in PC bilayers is stable, and can be used as the basis of more investigation.

For polarisable multipole simulations in AMOEBA, the inner and outer time steps of the r-RESPA integrator can be chosen independently. Preliminary simulations showed that stable integration was not possible with an inner time step of more than 1 fs (which is unsurprising given that hydrogen bonds and water model geometry are unconstrained in the AMOEBA model). The maximum stable outer time step was found to be 4 fs. Based on analysis of total energy, simulation box volume, and average temperature over the initial 0.25 ns of a 1 ns simulation with the AMOEBA forcefield (see **SI Fig. S2**). Based on this analysis, inner and outer time steps of 0.25 fs and 2 fs respectively were used for the production simulations. This is slightly more conservative than the time step of 2 used in the parameterisation of the AMOEBA protein force field [20]. Analysis of protein position relative to lipid bilayer after equilibration with the different force fields (data not shown) at the end of 10 ns long simulations using the additive CHARMM36 force field or the AMOEBA polarisable multipole force field showed similar positions of the M2 bundle in the bilayer for the various simulations. Taken together these results suggest that the system consisting of a pentameric M2 bundle in PC bilayers is stable, and so can be used as the basis of more detailed simulations and analysis.

#### *PMF convergence*

Convergence was assessed by calculating an additional PMF profile, $F_t(z)$, from a shorter time interval centered around time $t$, where an interval length of 2 ns was used. This instantaneous PMF profile can be compared to the final profile by means of the mean signed deviation, and the root mean squared deviation. For a converged PMF, the value of these error metrics tends to zero with increasing simulation time provided the interval length is large enough. See **SI Fig. S4**.

### SI Figures

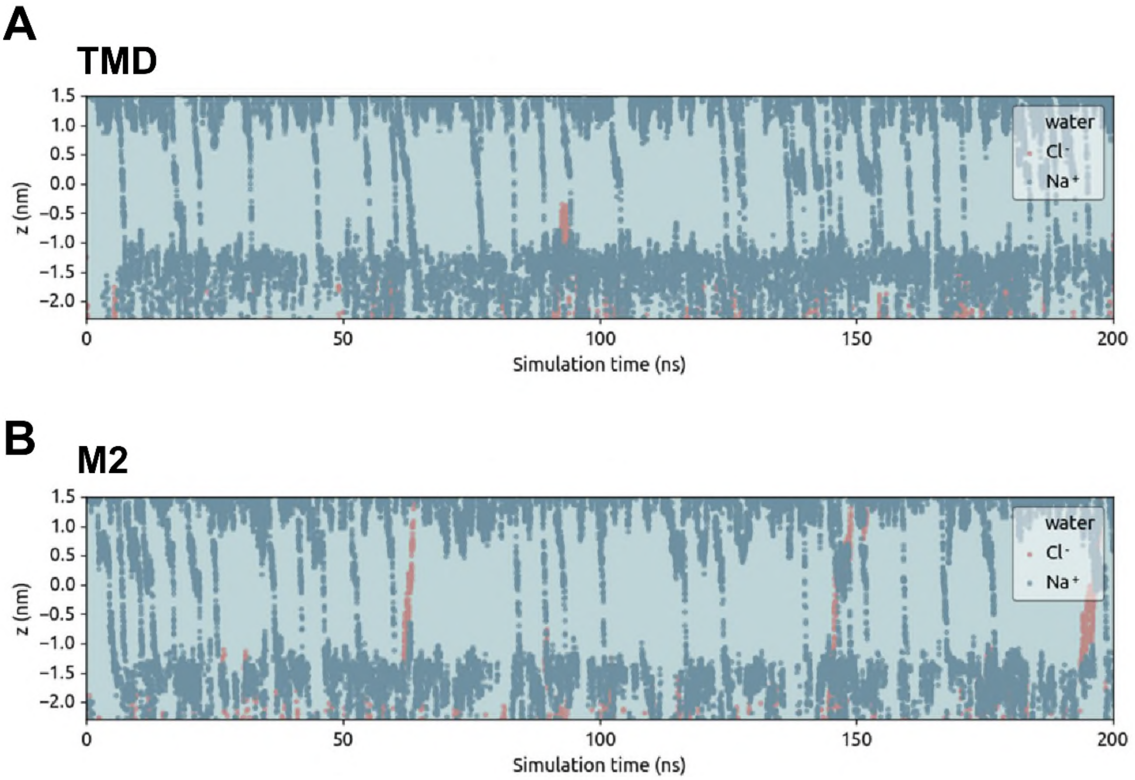


*SI Figure S1:*

Comparison of the 5HT3R M2 bundle vs. transmembrane domain in the presence of an applied electrostatic field (-500 mV, with negative potential on the cytoplasmic side). The graphs show ion and water trajectories through **A** the pentameric bundle of M2 helices and **B** the entire transmembrane domain (TMD) of the 5HT3R, both embedded in a phosphatidylcholine bilayer. Based on 3 x 200 ns of such simulations for each of the two systems, a similar level of conductance (corresponding to ~30 pS), with selectivity of sodium over chloride ions, was demonstrated.

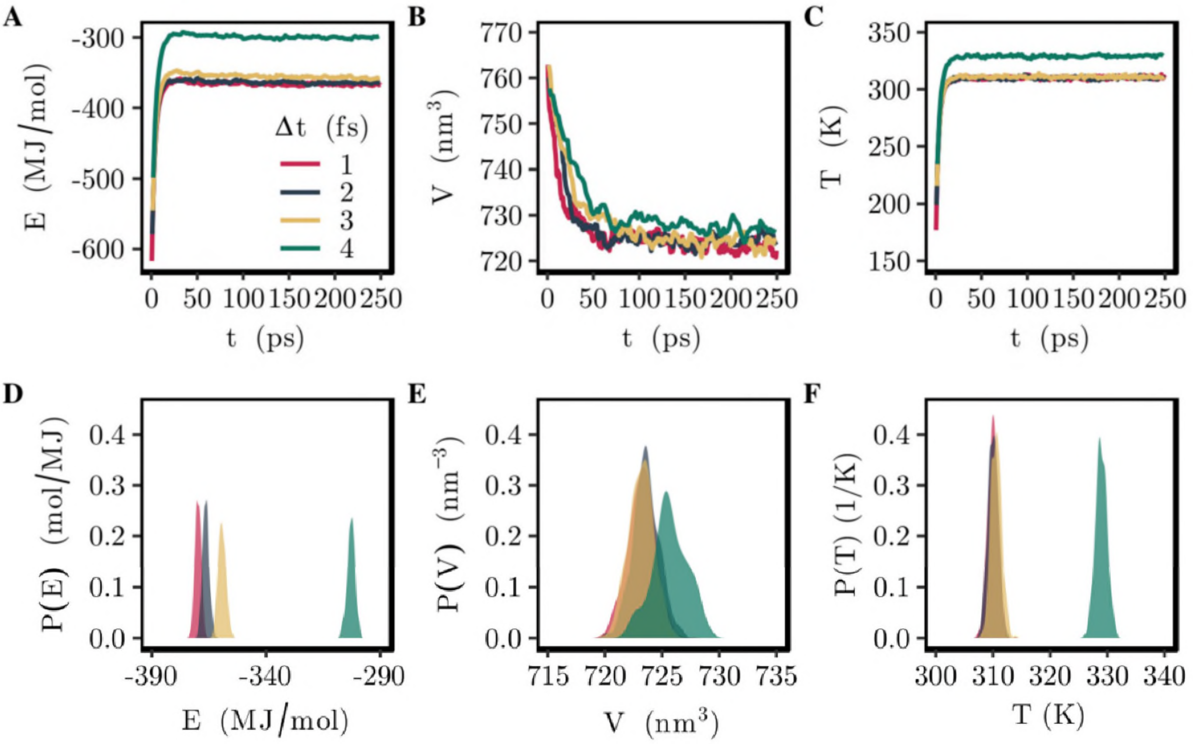


*SI Figure S2:*

Equilibration of the polarisable multipole simulation system. (**A – C**) Time series of total energy, E, box volume, V , and average temperature, T, over the initial 250 ps of a 1 ns long simulation with the AMOEBA force field. All three quantities converge to their equilibrium value within approximately 100 ps. (**D – F**) Probability densities estimated from samples taken from the equilibrated part of the trajectories (t > 250 ps). For an integrator time step of more than 2 fs, the thermodynamic properties of the system are inconsistent with those calculated using a more conservative time step. Data shown was derived from simulations employing the AMOEBA03 water model, and the closed state (PDB ID: 4PIR) of the pore.

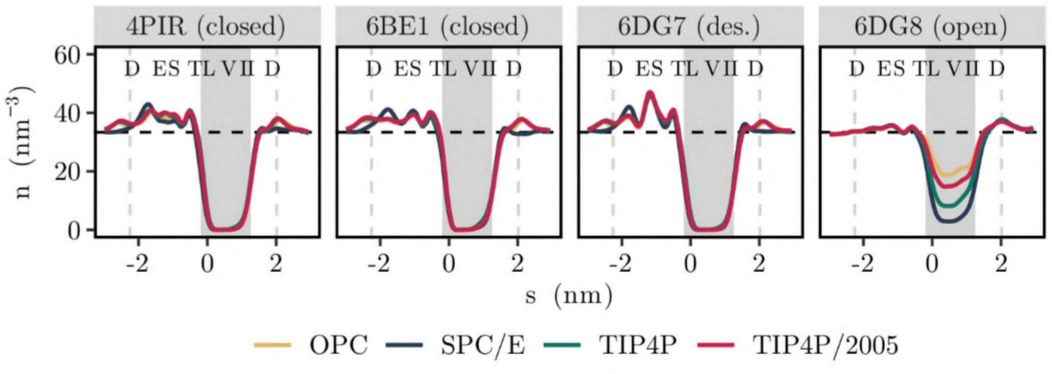


*SI Figure S3:*

Comparison of pore hydration using alternative additive water models. Water density in four conformational states of the 5HT3R. The profiles shown represent the average of three independent simulations (five in the case of the open state (PDB ID: 6DG8) structure) and the dashed horizontal line indicates the density of bulk water. A shaded background represents the hydrophobic gate region and dashed vertical lines denote the extent of the protein. Single letter codes for the pore-facing amino acid side chains are shown at the top of each panel.

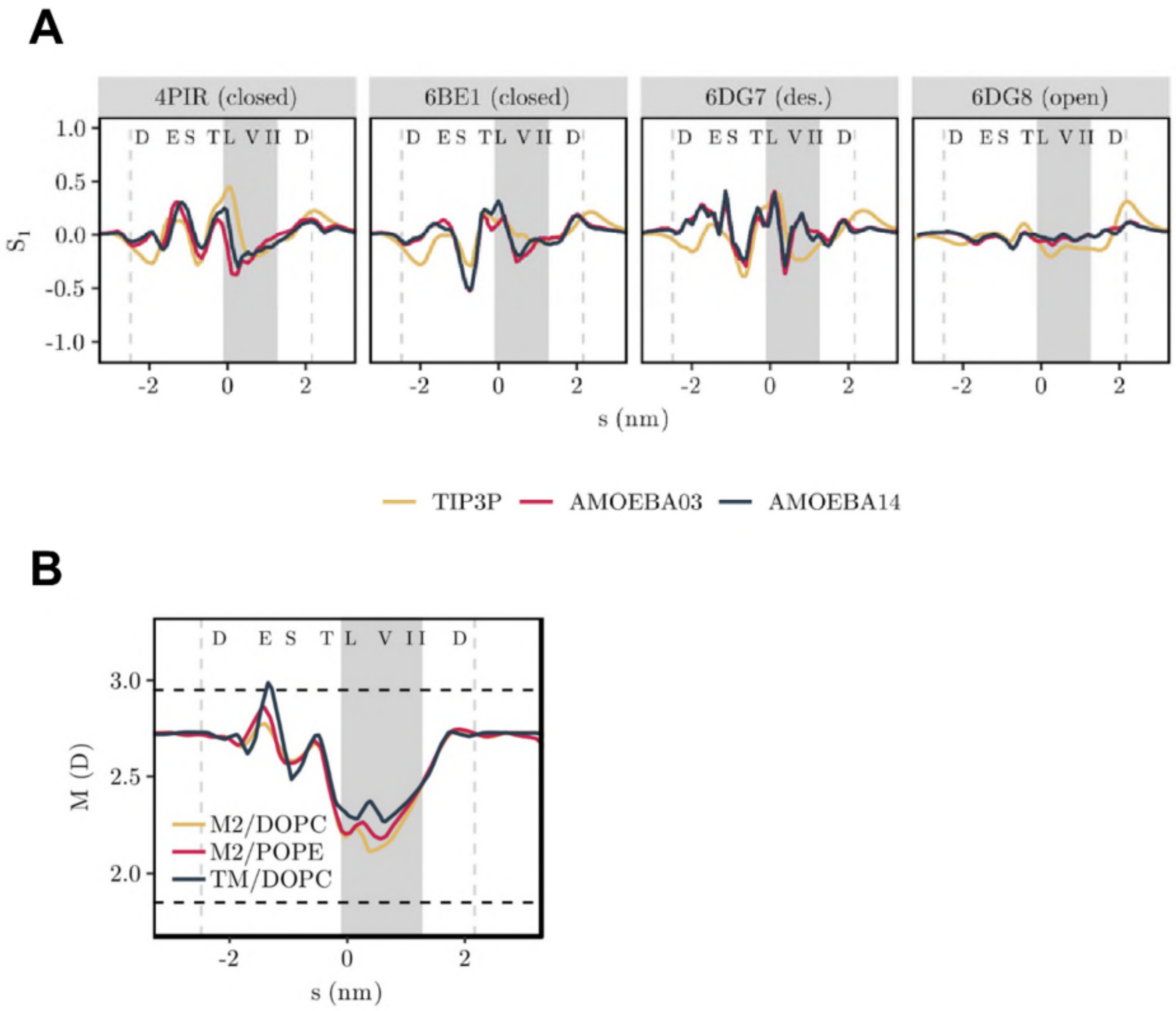


*SI Figure S4:*

**A** Molecular dipole moment of water in the pore of four conformational states of the 5HT3R showing the orientational order parameter of the molecular dipole moment, *S1*. The shaded area represents the hydrophobic gate region, in which de-wetting occurs. Dashed vertical lines indicate the extent of the simulated protein structure. Dashed horizontal lines at 1.85D and 2.95D represent the dipole moment of water in the gas and liquid phase respectively. Single letter codes for the pore-facing amino acid side chains are given at the top of each panel. Curves shown represent the average over three independent repeats. ***B*** Magnitude of the molecular dipole moment of water in three different simulation systems based on a closed state (4PIR) structure of the 5HT3R. Simulations employed the AMOEBA13 protein force field together with the AMOEBA14 water model. Curves shown represent the average over three independent repeats. The receptor was represented by either the M2 helix bundle embedded in a pure DOPC or pure POPE lipid bilayer or by the complete transmembrane domain embedded in a DOPC bilayer.

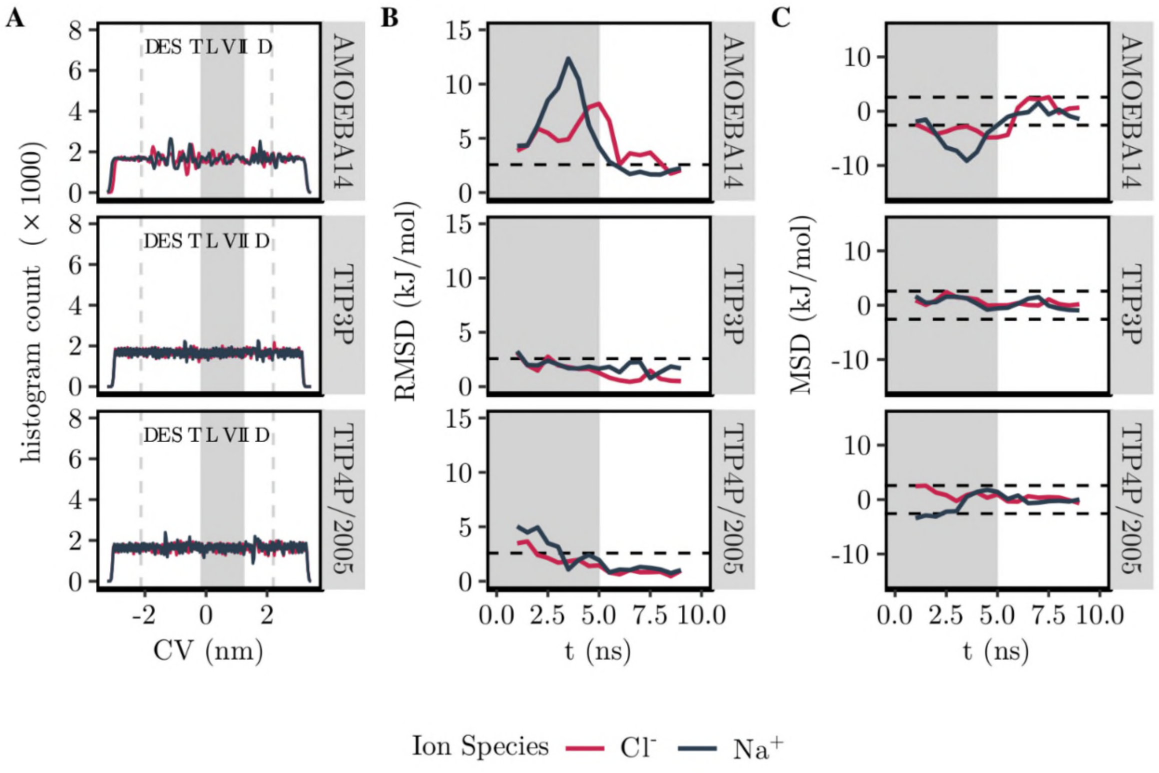


*SI Figure S5:*

Convergence of umbrella sampling simulations. (**A**) Histogram for both $Na^+$ and $Cl^-$ ions over all umbrella sampling windows. Dashed grey lines indicate the extent of the protein and the shaded area denotes the hydrophobic gate region. Single letter codes for the pore-facing amino acids are shown at the top of each panel. (**B,C**) Convergence of potential of mean force evaluated in terms of the root mean squared deviation (**B**) and mean signed deviation (**C**) with respect to the final profile. The shaded background indicates the 5 ns long equilibration period which is discarded in the calculation of the final PMFs shown in Figure 4 of the main text. Dashed horizontal lines indicate where the deviation falls below the thermal energy of 1 $k_BT$.

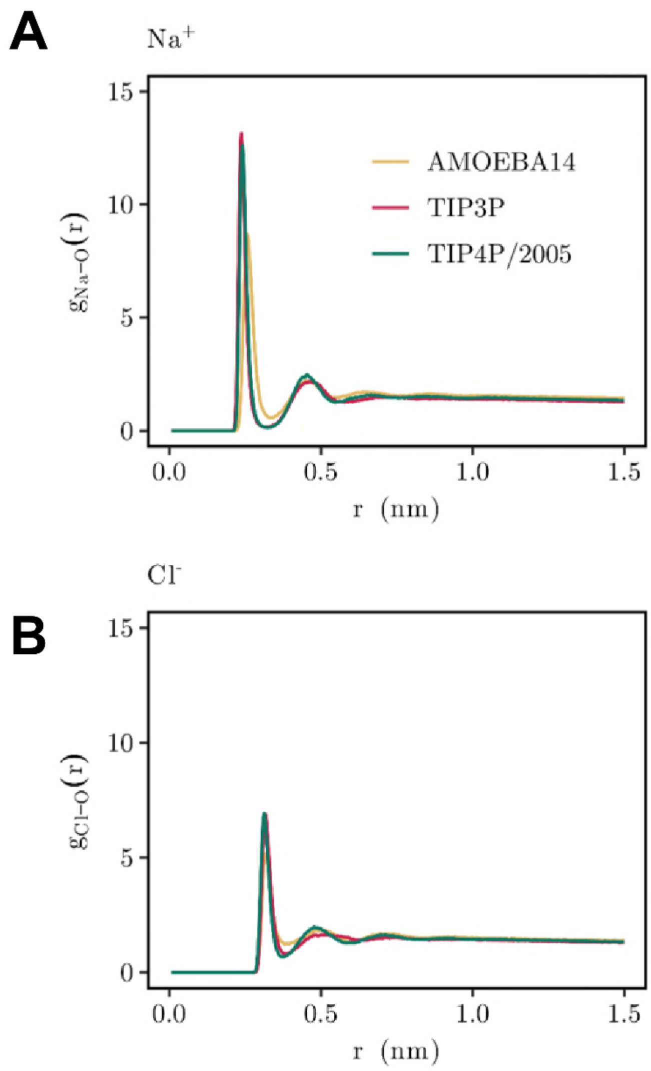


*SI Figure S6:*

Ionic hydration shells for different water models. Radial distribution function of water oxygens around $Na^{+}$ ions and $Cl^{-}$ ions in bulk water. The radial distribution function was calculated for an ion located in the bulk water regime outside the channel pore and is normalised to take a value of 1 at large separations.
